# High-Throughput Short Sequence Typing Schemes for *Pseudomonas aeruginosa* and *Stenotrophomonas maltophilia* pure culture and environmental DNA

**DOI:** 10.1101/2023.09.15.557936

**Authors:** Thibault Bourdin, Marie-Ève Benoit, Emilie Bédard, Michèle Prévost, Caroline Quach, Eric Déziel, Philippe Constant

**Author notes:** correspondence to Eric Déziel, and Philippe Constant,. **E-mail address of authors**: T. Bourdin; M-È. Benoit; E. Bédard; M. Prévost; C. Quach; E. Déziel; P. Constant.

## Abstract

Molecular typing techniques are employed to determine the genetic similarities between bacterial isolates. These methods primarily utilize specific genetic markers or analyze the complete genome sequence of pure bacterial cultures. However, the use of environmental DNA profiling to assess epidemiologic links between patients and their environment has not been explored in depth. This work reports on the development and validation of two High-Throughput Short Sequence Typing (HiSST) schemes targeting the opportunistic pathogens *Pseudomonas aeruginosa* and *Stenotrophomonas maltophilia*, along with a modified SM2I medium for specific isolation of *S. maltophilia*. Our HiSST schemes are based on four discriminative loci for each species and demonstrate high discrimination power, comparable to pairwise whole genomes comparison. Moreover, each scheme includes species-specific PCR primers, enabling precise differentiation from closely related taxa without the need for upstream culture-dependent methods. For example, the primers designed to target the *bvgS* locus allow to distinguish *P. aeruginosa* from the very closely related *Pseudomonas paraeruginosa* sp. nov. The selected loci included in the schemes for *P. aeruginosa* (*pheT*, *btuB*, *sdaA*, *bvgS*) and for *S. maltophilia* (*yvoA*, *glnG*, *ribA*, *tycC)*, are within the range of 271 to 330 base pairs adapted to massive parallel amplicon sequencing technology. A R-based script implemented in the DADA2 pipeline was assembled to facilitate HiSST analysis for efficient and accurate genotyping of *P. aeruginosa* and *S. maltophilia*. The performance of both schemes was demonstrated through *in-silico* validations, assessments against reference culture collections, and a case study involving environmental samples.

## Introduction

Molecular typing methods are invaluable tools in infection control, helping understand the source of specific strains and their relatedness in healthcare facilities, which is important for outbreak investigations. Primarily, molecular typing relies on multi-locus sequence typing (MLST) methods examining long DNA fragments from target genes (1), employing laborious procedures such as isolation of microorganisms by cultivation followed by Sanger sequencing of amplicons, or analyzing whole genome sequences (WGS) (2–5), and many other methods requiring pure cultures (6–10). However, the application of these methods to assess the epidemiological relatedness between patients and their environment through environmental DNA profiling has been relatively understudied. This knowledge gap might lead to an underestimation of pathogenic bacteria existing in a viable but non-culturable (VBNC) state (11, 12). While molecular typing has proven effective in characterizing strains and their transmission within healthcare environment, its utility in linking patients to environmental sources remains an underdeveloped opportunity. We have recently reported an innovative method called high-throughput short sequence typing (HiSST) to ease the monitoring of *Serratia marcescens* through PCR amplicon sequencing techniques (13). This novel approach improves the differentiation of bacterial isolates and facilitates the exploration of diversity profiles, even among non-cultured microbial populations through direct analysis of environmental DNA.

*Pseudomonas aeruginosa* is a Gram-negative rod-shaped bacterium, belonging to the Gammaproteobacteria clade, which is reported to be ubiquitous in environments impacted by human activities (14) and to thrive in moist and wet conditions (15). Due to its metabolic flexibility and high inherent resistance to antimicrobial agents, this species possesses the capacity to adapt to diverse ecological niches, encompassing hospital facilities and patient devices (8, 16–20). *P. aeruginosa* is a well-known opportunistic pathogen (21, 22), causing a wide variety of infections affecting most human organs (23). This species has garnered significant attention due to its clinical relevance and its widespread use as a model bacterium in various biological areas (24, 25). Consequently, a considerable number of genomes are available, including numerous clonal or closely related strains (https://pseudomonas.com/, https://ipcd.ibis.ulaval.ca/). This rich dataset enables comprehensive investigations and facilitates the development of new typing schemes, using next-generation sequencing such as HiSST. *Pseudomonas aeruginosa* is a Gram-negative rod-shaped bacterium, belonging to the Gammaproteobacteria clade, which is reported to be ubiquitous and to thrive in moist and wet conditions (15). Due to its metabolic flexibility and high inherent resistance to antimicrobial agents, this species possesses the capacity to adapt to diverse ecological niches, encompassing hospital facilities and patient devices (8, 16–20). *P. aeruginosa* is a well-known opportunistic pathogen (21, 22), causing a wide variety of infections affecting all human organs (23). This species has garnered significant attention due to its clinical relevance and its widespread use as a model bacterium in various biological areas (24). Consequently, a considerable number of genomes are available, including numerous clonal or closely related strains. This rich dataset enables comprehensive investigations and facilitates the development of new typing schemes, using next-generation sequencing such as HiSST.

*Stenotrophomonas maltophilia* was first classified in the genus *Pseudomonas* in 1961, then in the genus *Xanthomonas* in 1983, and since 1993 has been one of the more than fifteen species of the genus *Stenotrophomonas* (26). Known as an ubiquitous species in the environment, *S. maltophilia* can be found in aquatic and soil environments, in rhizospheres and on plants (27). This species is recognized as an opportunistic pathogen and one of the opportunistic premise plumbing pathogens associated with nosocomial infections (26, 28–30). Multiple selective media have been developed for *S. maltophilia* over the years (31–33), including the medium SM2I introduced by Adjidé in 2010 (34). However, its specific composition remains to be adjusted, and requiring a significant concentration of the costly antibiotic imipenem. To enhance the medium for *S. maltophilia* isolation, we report here a modified SM2I medium.

MLST schemes are already referenced and widely used for *P. aeruginosa* and *S. maltophilia* (35, 36). However, conventional MLST schemes are showing signs of becoming outdated due to their lack of primers specifically tailored to cover the entire range of strains, including more recent additions to databases. Moreover, these conventional schemes are not well-suited to large-scale epidemiological surveys due to labor-intensive upstream efforts in cultivation and isolation processes. This study was aimed at designing two HiSST schemes targeting the opportunistic pathogenic bacteria *P. aeruginosa* and *S. maltophilia*. The HiSST schemes were developed based on whole-genome sequences available in public databases, and validated with reference culture collections, environmental isolates and environmental DNA samples from neonatal intensive care units (NICUs). HiSST offers advantages in terms of accuracy, efficiency, and scalability in the context of large sample sizes, without the need for culture-dependent upstream procedures.

## Materials and Methods

### Development of the HiSST scheme

Pan-genome allele databases were assembled from 45 complete genomes of *P. aeruginosa* and 23 complete genomes of *S. maltophilia* retrieved from the NCBI GenBank database with the *Build_PGAdb* module available on PGAdb-builder online tool (37). Briefly, a preliminary step involved the identification of 37 and 40 highly conserved genes with the highest number of alleles for *P. aeruginosa* and *S. maltophilia*, respectively. After aligning the alleles of each gene and removing non-overlapping ends, a sequence identity matrix was computed using BioEdit (38) software. Gene fragments showing the highest polymorphism rate (i.e., highest nucleotide variations) and surrounded by conserved k-mers (allowing to design primers) were selected. Only target amplicon sequences under 350 bp size were chosen (length which is adapted for Illumina PE-250 sequencing) and subjected to alignment against the NCBI database using the Basic Local Alignment Search Tool (BLAST). This was carried out using a previously described step-by-step procedure to determine the specificity of the chosen gene fragments (13). Candidates for *P. aeruginosa* and *S. maltophilia* HiSST scheme included, respectively, 8 and 12 loci (i.e. nucleotide sequences of internal fragments of the previously selected genes) showing the highest polymorphism and the most specific non-overlapping ends. The selection procedure to achieve the identification of the four most discriminant loci was the same as described in our previous study (13).

### Primer design and PCR amplification

The primers used for both *P. aeruginosa* and *S. maltophilia* HiSST schemes were designed to specifically target internal loci as described above, composed of oligonucleotides that range in length from 17 to 22-mers (Table 1). To evaluate the specificity and coverage of the primers, we conducted *in-silico* tests using the software tool “Primer-BLAST” (39). The RefSeq non-redundant proteins database was used to assess the primers specificity, while the *P. aeruginosa* or *S. maltophilia* subset RefSeq database was used to verify the primers coverage for the corresponding species. The PCR reactions were carried out in 25 µL reaction volume containing 0.6 U Fast-Taq DNA polymerase (Bio Basic Inc., Markham, Canada), 1x of Fast-Taq Buffer (Bio Basic Inc., Markham, Canada), 200 µM dNTPs, 0.4 mg/mL Bovine Serum Albumin (Thermo Fisher Scientific Baltics UAB, Vilnius, Lithuania), 0.4 µM of each primer (except for *pheT* and *btuB* primers at 0.2 µM), and 2 ng/µL of extracted template DNA. A solution of 0.5x Band Sharpener (Bio Basic Inc., Markham, Canada) was included in all mixtures except for the *yvoA* locus of *S. maltophilia* HiSST scheme.

**Table 1:**
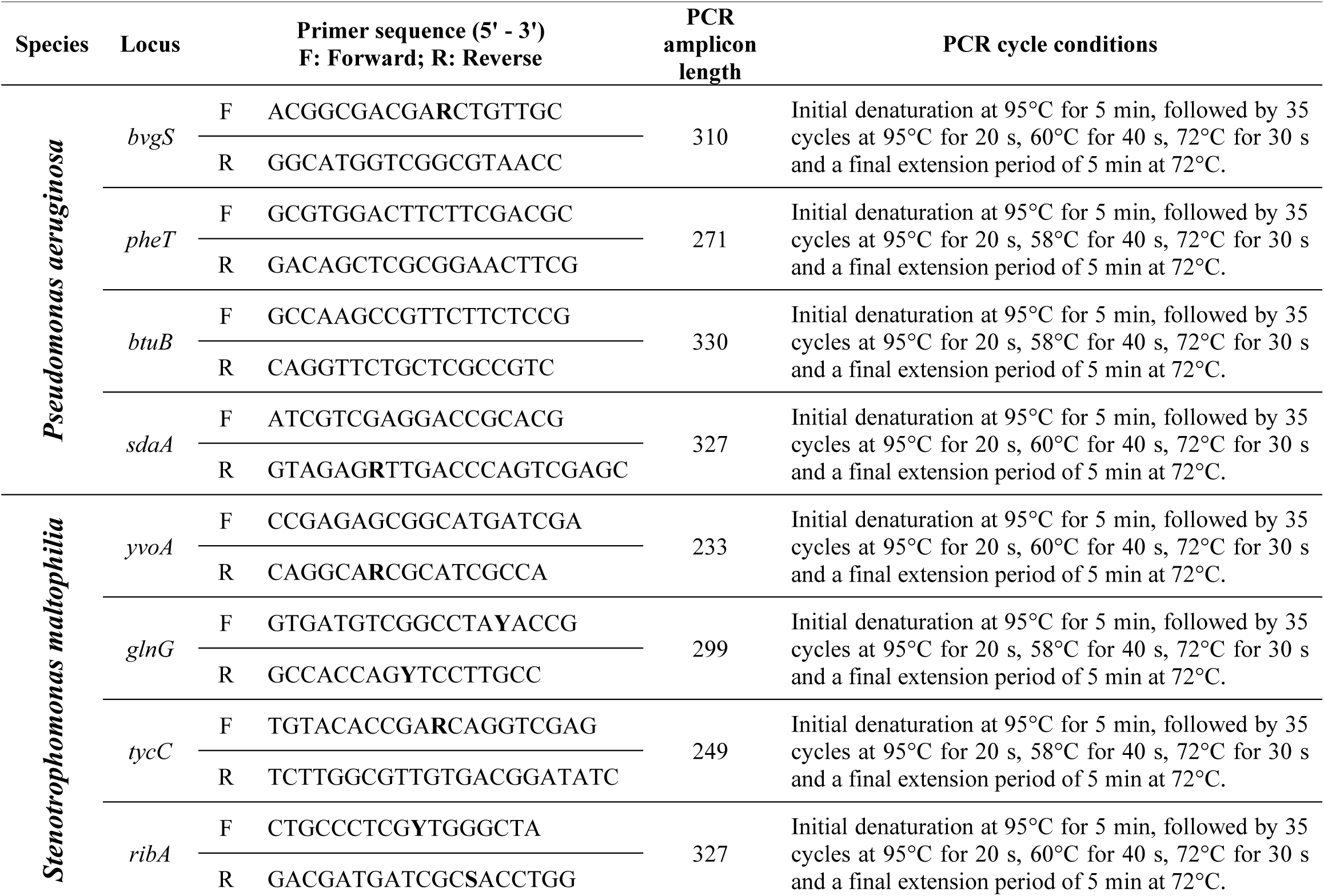
HiSST locus specific primers sequences and PCR cycle conditions*.

### Validation of the HiSST scheme with reference strains

Primers were validated *in-vitro* using reference strains obtained from diverse sources (Table S1). Selected strains for *P. aeruginosa* HiSST scheme comprised *P. aeruginosa* (n = 16), *P. paraeruginosa* (n = 1), *P. beteli* (n = 1), *S. maltophilia* (*n* = 2), *Stenotrophomonas acidaminiphila* (*n* = 1), *Serratia marcescens* (n = 2), *Serratia liquefaciens* (*n* = 1), *Serratia rubidaea* (*n* = 1), *Klebsiella pneumoniae* (*n* = 1), *Delftia tsuruhatensis* (*n* = 1), and *Staphylococcus haemolyticus* (n = 1). Our tests inadvertently included *S. haemolyticus* due to contamination of the *Pseudomonas* sp. storage culture we originally intended to examine (WGS was carried out to confirm the species). Primers from *S. maltophilia* HiSST scheme were tested for *S. maltophilia* (*n* = 16), *S. acidaminiphila* (*n* = 1), *Stenotrophomonas nitritireducens* (*n* = 1), *Stenotrophomonas rhizophila* (*n* = 1), *P. aeruginosa* (n = 1), *S. marcescens* (n = 2), *S. liquefaciens* (*n* = 1), *S. rubidaea* (*n* = 1), *K. pneumoniae* (*n* = 1), *D. tsuruhatensis* (*n* = 1). The strains were purified on Trypticase Soy Broth (TSB) (Difco Laboratories, Sparks, MD, USA – Le pont de Claix, France) with Agar (15 g/L) (Alpha Biosciences, Inc., Baltimore, MD, USA) at 30°C for 48 h. A single colony of each strain was inoculated in 2 mL TSB and grown for 48 h at 30°C for subsequent genomic DNA extraction.

### Validation of molecular typing by WGS

Three strains of *P. aeruginosa* and four strains of *S. maltophilia*, isolated from the environment, were subjected to WGS using the Illumina NextSeq 550 platform at SeqCenter (Pittsburgh, PA, USA). Trimmomatic v0.39 (40) was employed for Illumina adapter clipping and quality trimming, with a minimum average quality threshold of 30. Contigs assembly from the FASTQ files of paired– end reads was carried out using the SPAdes de novo assembler (41) and visualized using Bandage (42). The contigs obtained from the SPAdes output were aligned, ordered, and oriented using the most closely related reference genomes (*P. aeruginosa* PAO1 for *P. aeruginosa* genome assemblies and S*. maltophilia* NCTC10258 for S*. maltophilia* genome assemblies) through the utilization of the ABACAS tool (43) to generate a contiguous genome.

### ANI analyses

To illustrate the contrasting discriminatory power of the HiSST scheme and the whole genome sequences of *Pseudomonas* spp. or *Stenotrophomonas* spp., we employed a heatmap visualization to compute average nucleotide identity (ANI) values for both the selected HiSST loci and complete genomes. The estimation of percentage identities between strains was accomplished through ANIb (ANI based on BLAST+ alignment tool), using the Python package *pyani* (15). Then, heatmaps displaying ANIb values for the HiSST schemes and whole genome profiles were generated within RStudio environment (44) using *circlize* (45) and *ComplexHeatmap* (46) packages, and R scripts available on GitHub (https://github.com/TBourd/R_scripts_for_HiSST_scheme). The ANIb values were determined by combining the four loci from both HiSST schemes. If a locus was absent in a particular strain, the ANIb score was zero. This situation arose for negative controls *Pseudomonas* spp. or *Stenotrophomonas* spp. due to the use of highly specific primers, *ribA* and *bvgS*, which exclusively target *S. maltophilia* and *P. aeruginosa* species, respectively. It appears that other species do not possess these genes or the specific k-mers targeted by these primers.

### cgSNPs analyses

Core genome single nucleotide polymorphisms (cgSNPs) were identified for each strain using the Snippy pipeline v4.6.0 (47). The reference genomes used for mapping were *P. aeruginosa* PAO1 and *S. maltophilia* NCTC10258. The haplotype network tree was constructed using the SNiPlay pipeline v3 by uploading the VCF file (48) obtained through the *snippy-multi* script (49). The tree visualization was performed in R Studio using the *ape* package (50).

### SNP and HiSST profile analyses

The diversity of each sequence types included in *S. maltophilia* and *P. aeruginosa* HiSST profiles was visualized with minimum spanning trees (MST). The dataset type “Aligned Sequences (FASTA)” was employed by merging DNA sequences of the four loci (from the corresponding HiSST scheme), associated with a ST identifier, resulting in the creation of a single sequence per ST. Then, “geoBURST distance” algorithm was used to compute a full MST, using the PHYLOViZ software, version 2.0a (51). Finally, the eBURST program was employed to explore clonal complexes (52).

### Validation of the HiSST scheme with environmental samples

A few clinical samples were selected to demonstrate the practical application of the HiSST schemes under real conditions. The samples were collected in two NICUs located in Montreal (Québec, Canada). Sink drains and diapers worn by newborns were sampled, leading to three positive samples for *P. aeruginosa* and four positives for *S. maltophilia*. This selection included samples with distinct and identical HiSST profiles (= ST) originating from various sources (fecal matter from a newborn, sink drains, or faucet samples) and sample types (eDNA and isolates). Each sample was subjected to direct eDNA analysis and selective culture to assess the accuracy of HiSST analysis. The accuracy was evaluated by performing WGS on environmental or clinical isolates (13).

### Selective culture conditions for *P. aeruginosa*

To identify cultivable opportunistic pathogens, samples were inoculated on media selective for the species of interest. The presumptive isolates were confirmed by HiSST molecular analysis. Cetrimide medium (43.3 g/L Cetrimide Selective agar, 10 mL/L glycerol, supplemented with the antibiotic nalidixic acid at 15 µg/mL) was used for growth and selection of *P. aeruginosa* (53). The inoculated agar plates were incubated at the selective temperature of 42°C for 48 h, also promoting the production of pigmentation by *P. aeruginosa* (e.g., pyocyanin).

### Adaptation of a selective agar for *S. maltophilia*

To optimize the detection of *S. maltophilia* presence in diverse HiSST PCR-positive samples, we improved the SM2I medium by adjusting its composition and replacing the expensive antibiotic imipenem with a more cost-effective alternative, meropenem. Based on SM2I medium (34), modified SM2I (mSM2I) medium is composed of Mueller Hinton (40 g/L), maltose (40 g/L), DL-methionine (0.5 g/L) and bromothymol blue (0.06 g/L). The medium was adjusted to pH 7.1 before autoclaving, then supplemented with the antibiotics: vancomycin (6 mg/L), meropenem (16 mg/L), and the fungicide amphotericin B (4 mg/L). Inoculated mSM2I plates were incubated at 30°C for 48 h.

### Creating HiSST databases

The latest HiSST databases for the two HiSST schemes were updated in April 2023 (Table S2). The initial stage of database assembly involved gathering all short sequence types (SST) relevant to the HiSST scheme for the species under study. The nucleotide Basic Local Alignment Search Tool (BLASTn [51]) parameterized with the default settings was utilized to create the *P. aeruginosa* SST databases (Table S2.A), based on 513 genomes from The *Pseudomonas* Genome Database (https://www.pseudomonas.com/). *S. maltophilia* SST databases (Table S2.B) were generated by a BLASTn search on NCBI (https://blast.ncbi.nlm.nih.gov/Blast.cgi) conducted against the *S. maltophilia* group (taxid:995085), using nucleotide collection (nr/nt) and the “megablast” program optimized for highly similar sequences. Any sequences with unexpected alignment lengths were eliminated and only strains comprising the four selected loci were retained in specific databases to enable ST assignment of *P. aeruginosa* and *S. maltophilia* HiSST schemes (Table 1). The specific databases were utilized to create the SST database for each locus using a script within the RStudio environment (“Step3_Create_SST.R”, available in the GitHub repository: https://github.com/LaboPC/HiSST-schemes_TB). A ST identifier was generated for each HiSST profile using an R script (named “Step4_Assign_ST.R”), also accessible in the GitHub repository.

An unspecific database, comprising the closest relatives of targeted sequences, was created to include all nucleotide sequences targeted by HiSST primers, excluding the target species. This database is designed to facilitate taxonomic assignment and identification of non-specific SST, involving sequences from environmental samples or isolates.

Databases were built by conducting a BLAST search on each locus using NCBI. The search was performed by specifying a query subrange from 1 to the expected length of the locus, as indicated in Table 1. For *P. aeruginosa* SST databases, the first query excluded *P. aeruginosa* (taxid:287), while the second query targeted the closely-related *P. paraeruginosa* (taxid:2994495) group. In the case of *S. maltophilia* SST databases, the *S. maltophilia* group (taxid:995085) was excluded from the search set. For all SST databases, the “Somewhat similar sequences (BLASTn)” program was utilized, with a maximum of 1000 target sequences. Only aligned sequences with query coverage ranging from 99 to 100 were included in unspecific databases.

Both specific and unspecific SST databases were merged into a single fasta file per locus, with the removal of primer k-mers using the R script “Optional_Remove_primers.R” available in the GitHub repository (https://github.com/LaboPC/HiSST-schemes_TB). The R script “Step5_Merge_correcting_db_seq.R” was then employed in three consecutive steps: (i) nucleotide sequences below the expected sequence length were eliminated, (ii) a reverse complement operation was performed to ensure uniform nucleotide sequence orientation, (iii) sequence names were formatted and the unspecific SST databases were combined with the SST databases for *P. aeruginosa* or *S. maltophilia*. Finally, a BLASTn database was constructed for each locus using the SST databases, employing the “makeblastdb” application from the BLAST+ executables (55).

### Bioinformatical pipeline for HiSST analysis

The DADA2 pipeline (56) was adapted to the analysis of HiSST schemes, incorporating additional steps to enhance the accuracy of taxonomic assignments. The pipeline encompassed the following key procedures: raw sequencing reads processing included primer sequences removal with the software Cutadapt v. 2.10 (57), default parameters specified in the package dada2 v1.8.0 (58) including error correction, denoising, and paired ends merging, and additional steps for HiSST analysis.

To enhance specificity, the BLASTn algorithm was utilized to filter the DNA sequences obtained from the previous step. Non-specific ASVs, corresponding to unexpected nucleotide length or unrelated to the bacterial species under study, were identified and removed. A taxon table was generated, containing only the ASVs specific to the bacterial species of interest. Additionally, two tables were created for further analysis. The first table captured the presence or absence of SST across various samples and facilitated the construction of Jaccard dendrograms to assess similarity between samples. The second table associated each sample with its corresponding ASV and assigned ST through exact matching, providing valuable information for subsequent analysis and database maintenance. The entire process can be executed using the “FunHiSSTDada2.R” unified function from the dedicated GitHub repository (https://github.com/LaboPC/HiSST-schemes_TB), executed through the “Script_RUN_FunHiSSTDada2.R” R script.

### HiSST nomenclature and assignation

The following nomenclature is used to identify HiSST loci and HiSST profiles: a “locus short sequence type” number (locus-SST) is assigned for each ASV of the individual loci; and the combination of multilocus SST of the overall HiSST profile is defined by a “sequence type” number (ST). The HiSST scheme database and R scripts are available on GitHub at URL: https://github.com/LaboPC/HiSST-schemes_TB.

### Accession number(s)

Raw sequencing reads have been deposited in the Sequence Read Archive of the NCBI in the BioProject PRJNA1009139, as well as assembled genomes.

## Results

### Design of the HiSST scheme for *P. aeruginosa*

The design of the HiSST scheme for *P. aeruginosa* involved a successive selection process. Initially, a total of 5039 alleles were identified, from which 38 alleles were carefully chosen based on their profiles, with each allele having 40 to 44 unique profiles among 45 strains. The selection process resulted in 33 alleles being retained (Table S3). To further refine the scheme, eight potential alleles were selected for subsequent development stages, based on their high average nucleotide polymorphisms (i.e., low identity scores) at the corresponding locus and the presence of conserved k-mers suitable for primer design. Finally, the four loci retained for *P. aeruginosa* HiSST schemes include gene fragments of *bvgS* (Virulence sensor protein BvgS precursor [PA2583]), *pheT* (Phenylalanine--tRNA ligase beta subunit [PA2739]), *btuB* (Vitamin B12 transporter BtuB precursor [PA1271]), and *sdaA* (L-serine dehydratase 1 [PA2443]).

The HiSST scheme for *P. aeruginosa* has a minimum similarity threshold of 96% among its 196 STs (Fig. S1). The pairwise ANIb genomic similarity score was greater than 98% to distinguish the different *P. aeruginosa* strains (Fig. 1B). In comparison, using the four concatenated loci of the *P. aeruginosa* HiSST scheme yielded a 94% ANIb similarity score between *P. aeruginosa* strains. In contrast, sequences of other *Pseudomonas* spp. retrieved from NCBI genomic database exhibited less than 84% nucleotide identity with the STs of the *P. aeruginosa* HiSST scheme (Fig. 1A, B). The discrimination performance of the HiSST scheme was also demonstrated during primer-BLAST tests. Specificity of the primers was supported by all hits affiliated to *P. aeruginosa* but one sequence from the strain NCTC10783 identified as a *P. fluorescens* isolate. Actually, the genome sequence of strain NCTC10783 corresponds to *P. aeruginosa* (59) in genome comparison analysis (Fig. 1A, B), further supporting the specificity of the HiSST scheme.

**Figure 1:**
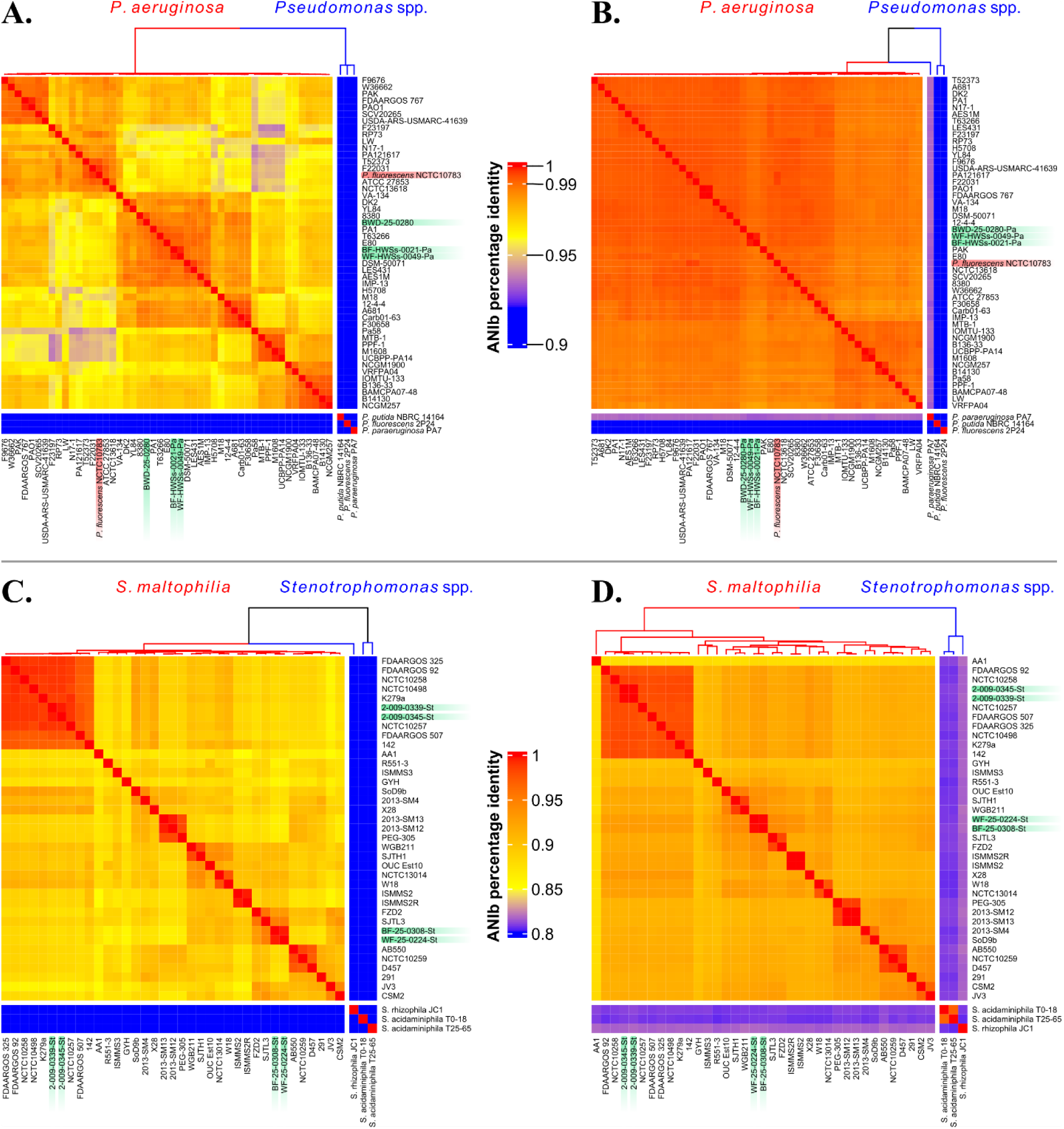
Discrimination of *Stenotrophomonas* spp. and *Pseudomonas* spp., based on the HiSST scheme and whole genome sequences. The heat-map reports the ANIb score of (A, C) the four concatenated loci of the HiSST scheme and (B, D) genome similarity. No *Stenotrophomonas maltophilia* and *Pseudomonas aeruginosa* were included as outgroup. The more cells turn from blue to red, the higher the ANIb score, and the more the strains are genotypically similar. Strains highlighted in green were derived from eDNA (BWD-25-0280) or DNA isolated (samples ending with “Pa” or “St”) from the NICU surveyed in this study. The strain highlighted in red is a misassignment of a *P. aeruginosa* strain as *P. fluorescens* (NCTC10783).

Each locus contained 17 to 36 informative sites (Table S4). STs profiles remained highly heterogeneous and distinguishable (Fig. 2.A). Among the 196 STs identified, the eBURST program classified eight potential clonal complexes (Fig. S1). Each group consists of only two STs, including two closely related groups (groups 6 and 7 at eBURST level 2). The diversity observed among *P. aeruginosa* strains was found to be representative of the whole-genome profiles (Fig. S2). Notably, clonal complexes were preserved, whether the analysis is based on whole-genome profiles or the HiSST scheme. Furthermore, strains sharing identical STs were found to be likely of clonal origin.

**Figure 2:**
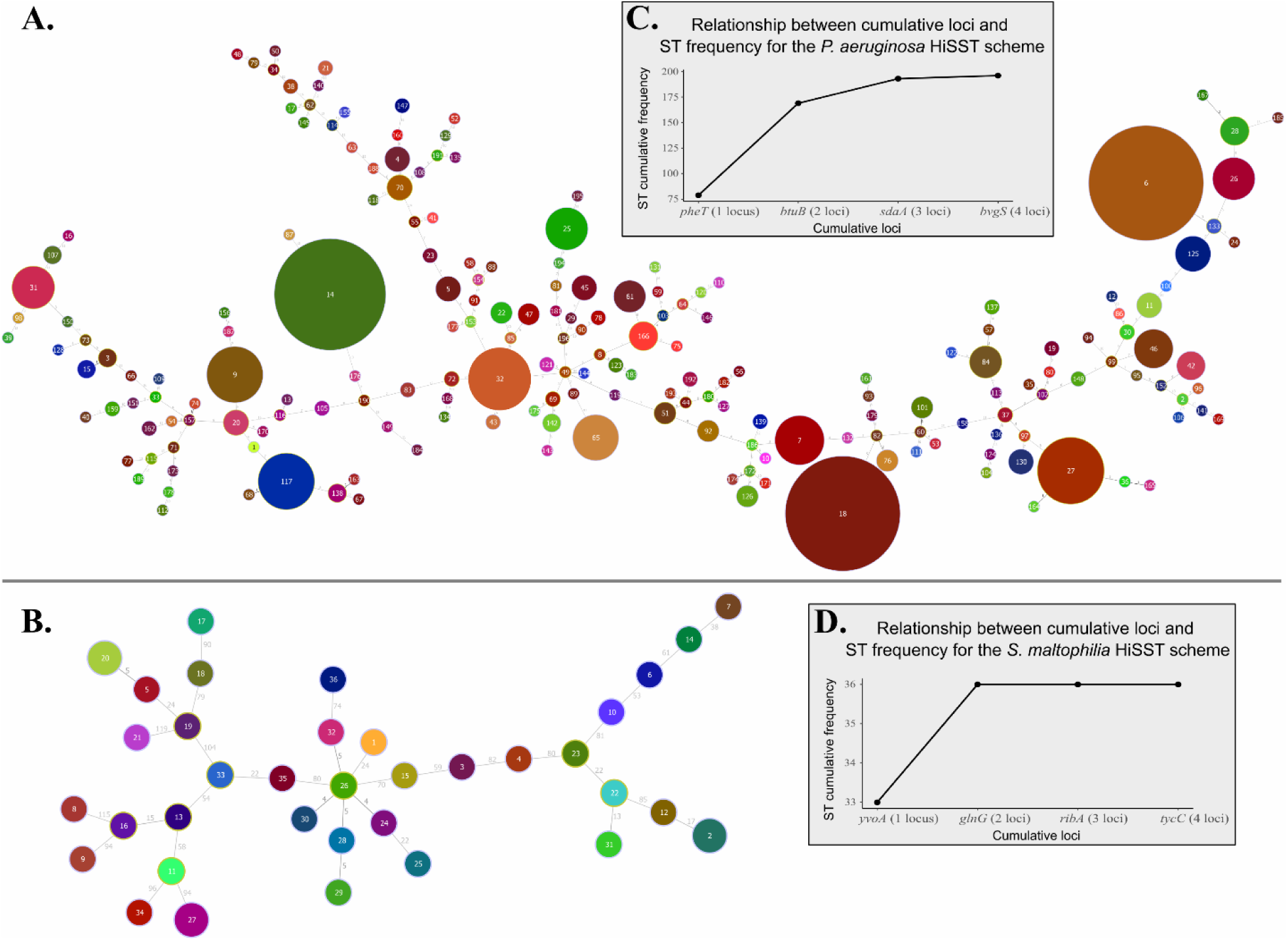
Minimum spanning trees for HiSST schemes of (A) *Pseudomonas aeruginosa* and (B) *Stenotrophomonas maltophilia*, based on SNP analysis. The distance labels represent the number of discriminating SNPs between neighbouring genotypes. Each pie chart label refers to sequence type (ST) identifier of the corresponding HiSST scheme. The ST diversity is displayed by the grey boxes, illustrating the relationship between ST frequency and the number of loci included in the (C) *P. aeruginosa* or (D) *S. maltophilia* HiSST scheme.

The number of selected loci (n = 4) for the HiSST scheme is deemed optimal for discriminating *P. aeruginosa* strains (Fig. 2C). In fact, the maximum cumulative frequency of STs was achieved using only three loci: *pheT*, *btuB* and *sdaA*. Among these loci, *pheT* demonstrated the highest discriminatory power, distinguishing the majority of *P. aeruginosa* strains, with 79 unique SSTs identified among 513 *P. aeruginosa* isolates (Table S2.A). This was followed by *btuB* (64 SSTs), *sdaA* (54 SSTs), and *bvgS* (35 SSTs). Although a notable decrease in discriminatory power was observed with the fourth locus, *bvgS*, it remains highly relevant for distinguishing *P. aeruginosa* from other *Pseudomonas* spp. Recently, Rudra et al. proposed a revised classification for the outlier clade of *P. aeruginosa* PA7 (= NCTC 13628T = ATCC 9027) as a novel species, *Pseudomonas paraeruginosa* sp. nov. (60). The HiSST scheme developed in this study is exclusively specific to *P. aeruginosa* due to the inclusion of locus *bvgS*, which is absent in *P. paraeruginosa*, as supported by BLASTn results and *in-vitro* tests conducted on *P. (para)aeruginosa* PA7 (= NCTC 13628T = ATCC 9027).

### Design of the HiSST scheme for *S. maltophilia*

For *S. maltophilia*, a total of 2077 alleles were initially identified, and from these, 40 alleles were selected based on their profiles, with each allele exhibiting 22 unique profiles across 23 strains. This selection process resulted in 39 alleles being retained (Table S3). Among these, eight potential alleles were chosen for further development of the HiSST scheme. Ultimately, four gene fragments were retained, encompassing *yvoA* (HTH-type transcriptional repressor YvoA), *glnG* (Nitrogen assimilation regulatory protein), *tycC* (Tyrocidine synthase 3), and *ribA* (GTP cyclohydrolase-2). The minimal similarity threshold among *S. maltophilia* strains was determined to be 86% for the four loci (Fig. S3). The ANIb similarity score for the HiSST scheme was determined to be 85%, effectively distinguishing between *S. maltophilia* strains. In comparison, the ANIb score obtained through whole-genome analysis is over 87% (Fig. 1C, D). Furthermore, genomes that belong to the same ST are undoubtedly clones, as also demonstrated by the cgSNP analysis (Fig. S4). These results suggest a comparable discriminatory power between whole-genome analysis and the HiSST scheme.

In contrast with *P. aeruginosa*, the loci targeted by the *S. maltophilia* HiSST scheme exhibited a high diversity rate, with a range of around 10% estimated diversity for the four loci, encompassing 62 to 81 informative sites (Table S4). This contributes to the high heterogeneity of STs, even though the database for *S. maltophilia* is not as extensive as that of *P. aeruginosa*, with only 39 complete genomes available compared to 513 strains used for *P. aeruginosa* HiSST scheme. No clonal complex was identified using the eBURST program.

Among the four loci, *yvoA* and *glnG* demonstrated the highest discriminatory power, with 33 unique SSTs identified, followed by *ribA* (31 SSTs) and *tycC* (30 SSTs). The maximum cumulative frequency of STs was achieved using only *yvoA* and *glnG* (Fig. 2D.), but *ribA* demonstrated the highest species-specificity to *S. maltophilia*.

### mSM2I agar selective for *S. maltophilia*

mSM2I was found to be highly specific for the selection of *S. maltophilia*, effectively improving its detection in various environmental samples (Bourdin et al., in preparation). All the reference strains tested for the *S. maltophilia* HiSST scheme (Table S1) were also grown on mSM2I. The medium successfully inhibited the growth of other species, including strains from three other *Stenotrophomonas* species, while allowing *S. maltophilia* to thrive after 48 h of incubation at 30°C. The medium mSM2I appears green at neutral pH, but turns blue in the presence of *S. maltophilia* as the pH rises into the alkaline range. *S. maltophilia* colonies are easily recognized by their olive-green to dark hue.

### Validation and Application of the HiSST Schemes

The primers, originally designed in 2019, have maintained their specificity up to time of writing this report (May 2023), as confirmed by a recent *in-silico* analysis, and subsequent laboratory validation through PCR testing using reference strains (Fig. S5 and S6 represent tests conducted in 2023). All *P. aeruginosa* and *S. maltophilia* strains exhibited optimal PCR amplicon of the correct size when subjected to their respective HiSST primers. As anticipated, amplification was observed for *P. paraeruginosa* with primers *pheT*, *btuB* and *sdaA*, while no amplification was detected when using primers designed to target the *bvgS* locus, thereby indicating a higher specificity for *P. aeruginosa*. None of the four primer pairs yielded amplification for other *Pseudomonas* spp. or other bacterial genera tested.

The *ribA* primer pair displayed the utmost specificity of the HiSST scheme of *S. maltophilia*, resulting in amplification exclusively for the targeted species. However, amplifications were observed for *S. acidaminiphila* and *S. nitritireducens* with the *tycC*, *glnG* and *yvoA* primers. Additionally, the *glnG* primer demonstrated the ability to amplify each tested *Stenotrophomonas* spp. Although certain primers lack species specificity, these limitations are mitigated by the inclusion of highly specific primer pairs (*bvgS* for *P. aeruginosa* and *ribA* for *S. maltophilia*), the genus-level specificity of all primers, and the integration of BLAST analysis into the DADA2 pipeline, thereby enhancing the analysis of HiSST schemes and facilitating the removal of unspecific ASVs, if necessary.

The primers have been utilized in an extensive study aimed at characterizing the ecology of three opportunistic pathogens, including *P. aeruginosa* and *S. maltophilia*, in the sink environment of two NICUs, resulting in the HiSST analysis of thousands of samples (Bourdin et al., in preparation). The results supported the specificity of each primer sets, even when applied to complex samples containing a high concentration and diversity of bacteria, such as biofilms or water from sinks P-traps. There was a concordance between HiSST schemes and ANIb analysis of whole genomes (Fig. 1), as well as cgSNPs (Fig. S4). Regarding *P. aeruginosa*, two faucet samples (aerator and tap water) collected from the same sink over a one-month time interval exhibited the same ST (samples BF-HWSs-0021-Pa and WF-HWSs-0049-Pa). Strains were successfully isolated from both samples, genotyped by HiSST and analyzed by WGS, confirming them as likely clonal strains (Fig. 1B and in green on Fig. S2). Another eDNA sample, BWD-25-0280, obtained from biofilm and water in a P-trap, also tested positive for *P. aeruginosa*. The strain was successfully isolated and exhibited the same HiSST profile as the eDNA sample, which was further confirmed by WGS.

In the investigation of *S. maltophilia*, two faucet samples (aerators and tap water) were found to be colonized by the same ST, as indicated by HiSST analysis of eDNA samples. HiSST analysis of *S. maltophilia* isolates confirmed their identical genotypes, which were further validated by WGS, confirming their clonality. A similar case was observed in fecal samples collected from the same newborn at different dates (one week apart), where the same ST was identified in both samples. The isolation of strains and subsequent WGS analysis confirmed the consistency of these findings.

## Discussion

Identifying discriminant alleles for *P. aeruginosa* poses a significant challenge. Strain differentiation primarily arises from the presence or absence of genes rather than nucleotide polymorphisms, indicating extensive genomic plasticity largely attributed to frequent horizontal gene transfers (25, 61, 62). HiSST is a robust genotyping tool that provides highly reliable predictions for assessing clonality between strains. However, in cases where two strains exhibit identical HiSST profiles, it is highly recommended to employ WGS, especially for epidemiological investigations involving *P. aeruginosa* (63). WGS analysis can confirm if two isolates are clonal, despite the low probability of non-clonal strains having the same HiSST profile within a single sample. Like for any typing methodologies, it is important to note that the presented HiSST scheme, while effective for the current strain repertoire, may eventually need to be updated in the context of the evolving database and the inclusion of new strains that might not be targeted by the existing HiSST primers. This will ensure its continued applicability and accuracy in capturing the full diversity of *P. aeruginosa* strains. An intriguing finding of this study is the exclusive presence of the *bvgS* allele in *P. aeruginosa* and its absence in all other *Pseudomonas* species. This distinctive genetic marker further strengthens the ability to differentiate *P. aeruginosa* from related *Pseudomonas* species, which include the closely related *Pseudomonas paraeruginosa* sp. nov, underlining the specificity and utility of the HiSST scheme.

The loci in the *S. maltophilia* HiSST scheme demonstrated a higher level of polymorphism, than those of *P. aeruginosa*, contributing to the heterogeneity of STs. While the database for *S. maltophilia* is limited compared to *P. aeruginosa*, the scheme showed promising specificity, with the *ribA* locus being highly specific to *S. maltophilia*. Indeed, the *ribA* locus is seldom found in other *Stenotrophomonas* spp. Furthermore, the primers designed for *ribA* are exclusively specific to *S. maltophilia* (Fig. S6 and primer-BLASTn query). Including the fourth locus *tycC* takes a more conservative approach to ensure maximal genotyping distinction. This consideration takes into account the limited size of the *S. maltophilia* database and the eventual occurrence of unknown genotype variants where certain loci may fail to amplify.

Both HiSST schemes demonstrate superior or similar differentiating power for *P. aeruginosa* and *S. maltophilia* strains compared to the ANI score, based on whole-genome comparisons, and the cgSNPs analysis. HiSST schemes effectively reflect the clustering of strains according to their whole genomes, enabling the discrimination of strains and identification of potential clonal strains. Furthermore, the analysis of HiSST schemes was enhanced by incorporating additional steps and adaptations into the DADA2 pipeline. This improvement led to more accurate taxonomic assignment and the generation of informative tables, facilitating in-depth investigations and analysis.

In conclusion, we developed HiSST schemes for *P. aeruginosa* and *S. maltophilia* based on four loci of conserved, yet polymorphic, genes. As a demonstration, we utilized the HiSST schemes on samples collected from a hospital environment to successfully achieve the characterization of genotype diversity of *P. aeruginosa* and *S. maltophilia*. The results demonstrate the accuracy and reliability of the HiSST genotyping method, both in comparison to whole-genome ANIb and cgSNPs analysis, and through the confirmation of clonal relationships using isolates and WGS. The genotyping schemes developed here should represent powerful tools for both ecological and epidemiological investigations. They enable the detection and identification of bacterial genotypes from environmental or isolated DNA in various samples, addressing limitations of culture-dependent methods that might underestimate bacteria, including pathogens present in a viable but non-culturable (VBNC) state within biofilms (11, 64).

## Acknowledgments

We thank the hospital staff for their help with sampling, Ann Brassinga (Department of Microbiology, University of Manitoba), Jonathan J. Ewbank (Centre d’Immunologie de Marseille-Luminy, Aix-Marseille University, Marseille, France), and the Laboratoire de santé publique du Québec for providing reference strains.

This work was supported by NSERC and CIHR through the Industrial Chair on Drinking Water and the Collaborative Health Research Program funding (CHRP 523790-18). Dr. Emilie Bédard is supported through a salary award from the Fonds de recherche du Québec – Santé (Junior 1); Dr. Caroline Quach is the Canada Research Chair (Tier 1) in Infection Prevention.

